# *adgrl3.1*-deficient zebrafish show noradrenaline-mediated externalizing behaviors, and altered expression of externalizing disorder-candidate genes, suggesting functional targets for treatment

**DOI:** 10.1101/2023.01.25.525492

**Authors:** Barbara D. Fontana, Florian Reichmann, Ceinwen A. Tilley, Perrine Lavlou, Alena Shkumatava, Nancy Alnassar, Courtney Hillman, Karl Ægir Karlsson, William H.J. Norton, Matthew O. Parker

## Abstract

Externalising disorders (ED) are a cause of concern for public health, and their high heritability make genetic risk factors a priority for research. Adhesion G Protein-Coupled Receptor L3 (*ADGRL3*) is strongly linked to several EDs, and loss-of-function models have shown impacts of this gene on several core ED-related behaviors. For example, *adgrl3.1^-/-^* zebrafish show high levels of hyperactivity. However, our understanding of the mechanisms by which this gene influences behavior is incomplete. Here we characterized, for the first time, externalizing behavioral phenotypes of *adgrl3.1^-/-^* zebrafish and found them to be highly impulsive, show boldness in a novel environment, have attentional deficits, and show high levels of hyperactivity. All of these phenotypes were rescued by atomoxetine, demonstrating noradrenergic mediation of the externalizing effects of *adgrl3.1*. Transcriptomic analyses of the brains of *adgrl3.1^-/-^* vs wild type fish revealed several differentially expressed genes and enriched gene clusters that were independent of noradrenergic manipulation. This suggests new putative functional pathways underlying ED-related behaviors, and potential targets for the treatment of ED.

## Introduction

The global public health and economic burden associated with untreated or unaddressed externalising behaviors (alcohol and substance misuse, violence, and aggression, oppositional or disruptive behavior) are significant. Externalising disorders (ED), such as attention-deficit/hyperactivity disorder (ADHD), conduct disorder, oppositional defiant disorder, or substance use disorder, are common among young people: ADHD ~4%^1^; Substance Use Disorders ~4%; conduct disorders ~3%^1^. ED are characterized by several transdiagnostic phenotypes, including inattention, hyperactivity and poor impulse control ^2–4^. As well as disrupting development and education, ED are associated with a range of debilitating comorbidities (bipolar disorder^5^, depression^6, 7^, anxiety ^8–10^, substance misuse ^11, 12^and sleep disorders^13^).

The high degree of heritability of ED (~80%^14^) has motivated the search for candidate genes. Recent linkage studies have identified several variants in the *ADHESION G-PROTEIN COUPLED RECEPTOR L3* (*ADGRL3*) gene that increase susceptibility to, and severity of, several EDs. For example, *ADGRL3* is linked to an increased risk of ADHD diagnosis and its clinical manifestation, and affects the efficacy of psychostimulant treatments ^15^. *ADGRL3* variants are also linked to substance use disorders ^12^. Several *ADGRL3* variants are associated with the clinical progression of ADHD and may have a similar impact on other externalising behaviors. For example, *ADGRL3* variants cause a 6-fold increase in risk for ADHD^16^, increase the persistence of combined-type ADHD into adulthood ^17^, increase impulse control problems in ADHD patients ^18^, and increase symptom severity in ADHD patients^19–21^. Despite being strongly associated with ADHD and several other impulse-control disorders (*e.g.*, substance abuse)^12,15^ the underlying mechanisms by which *ADGRL3* affects externalising behavioral phenotypes are not well characterized. Our aim in this study was to perform a behavioral and molecular characterisation of *adgrl3* to understand its role in promoting or mediating externalising behaviors.

*ADGRL3* codes for a G-protein coupled receptor involved in cell adhesion, signal transduction and synaptic signalling^19, 22, 23^. It is widely expressed throughout the brain, including the prefrontal cortex (PFC), caudate nucleus, amygdala and cerebellum, and across a diverse range of CNS structures and nuclei including the cerebrum (frontal and temporal lobes, occipital pole), limbic system (hippocampus) and striatum (putamen^22^. *ADGRL3* is conserved across a range of taxa^24–26^, and knocking out or knocking down this gene causes a range of ADHD-relevant symptoms. For example, *Adgrl3^-/-^* rats and mice, and both *adgrl3.1* antisense morpholino oligonucleotide (MO) knock-down and *adgrl3.1^-/-^* knock-out zebrafish larvae display persistent hyperactivity^24, 27^ which is rescued by both stimulant and non-stimulant ADHD treatments (e.g. the dopamine (DAT) reuptake inhibitor methylphenidate^28^ and the noradrenaline (NET) reuptake inhibitor atomoxetine^29 28 24^.

As well as hyperactivity, *Adgrl3^-/-^* mice display decreased inhibitory control^26^. Inhibitory control is a multifaceted process that includes the inhibition of prepotent responses, intolerance of delay, and preference for small-immediate vs large-delayed rewards^3^. Despite being dissociated from one another both clinically and neurologically, and impulse control-subtypes rarely inter-correlating at the interindividual level^30^, deficits in all impulse-control subtypes are reversed by the selective noradrenergic reuptake inhibitor atomoxetine, suggesting a general mechanistic role for noradrenaline in this process^31^. In addition, extensive previous research has demonstrated that atomoxetine significantly reduces impulse control deficits in rodents^31^, humans^32^ and zebrafish^33^. In terms of hyperactivity, we recently carried out a screen of *adgrl3.1^-/-^* zebrafish larvae and found several drugs rescue hyperactivity, including aceclofenac, amlodipine, doxazosin, and moxonidine^29^. Although the molecular mechanisms of action by which this gene impacts externalizing behaviors is not clear, collectively, these data suggest that *ADGRL3* is functionally involved with two core externalising symptoms (hyperactivity and reduced impulse-control) representing an ideal tool for identifying targets for the development of novel therapeutics.

In order to characterise the contribution of *adgrl3.1* to externalising symptoms such as poor impulse control, inattention and hyperactivity, we first characterized these behavioral phenotypes in *adgrl3.1^-/-^* zebrafish, and validated this with atomoxetine treatment (commonly used as treatment for externalizing disorders). We found that *adgrl3.1^-/-^* show marked deficits in impulse-control and attention and increased hyperactivity, all of which are rescued by atomoxetine. We then carried out a fine-grain examination of the brain transcriptome of *adgrl3.1^-/-^* compared to wildtype controls, to characterise alterations in gene networks and biological pathways. In addition, we examined immediate changes in gene expression that occur when *adgrl3.1^-/-^* are treated with atomoxetine. We identified several differentially expressed genes (DEGs) between WT and *adgrl3.1^-/-^* including *dusp6,* offering insights into the mechanism by which *ADGRL3* may mediate externalizing behaviors, and suggesting potential targets for treatment.

## Materials and Methods

### Generation of *adglr3.1^-/-^* fish

The fish used in this study were homozygous *adgrl3.1* mutant zebrafish (*adgrl3.1-/-*) created by CRISPR-Cas9 genome engineering, as previously described^29^, and age-matched *adgrl3.1*+/+ controls. Adult fish were housed in an aquarium facility at the University of Portsmouth (UK), and kept on a 14:10 light: dark cycle (28°C, pH~8). *adgrl3.1^-/-^* and controls (wild-type AB strain) were simultaneously in-crossed (pair breeding) and grown on a recirculating rack (Aquaneering, USA) to 4 months-post fertilisation before testing. All housing tanks contained enrichment substrates from 10-days post fertilisation and throughout (gravel pictures under the tanks). Offspring from *adgrl3.1^-/-^* and wild-type controls were randomly selected from ~5-10 groups of 20, with equal numbers of males and females in each experimental group (detailed below). All work was carried out following scrutiny from the University of Portsmouth Animal Welfare and Ethical Review Body (AWERB), and under licence from the UK Home Office (PPL P9D87106F).

### Drug treatment

*adgrl3.1^-/-^* fish were individually treated with 0.5 mg/L atomoxetine (TCI UK Ltd., Oxford, UK) for 30 minutes prior to behavioral recording. The concentration used was based on extensive previous research from our group and others^33^. The drug was dissolved in aquarium-treated water and animals were individually treated in 300 mL beakers. For the 5-CSRTT, atomoxetine treatment followed establishment of steady state responses on the final phase of the 5-CSRTT.

### RNA sequencing analysis

Fish were euthanized by rapid cooling (immersion in 2°C water) and the whole brain tissue was removed, snap-frozen in liquid nitrogen, and kept at −80°C until further use. RNA extraction was performed using the GeneJET RNA Purification Kit (Thermo Scientific) as described in the manufacturer’s instructions. Next, RNA concentration was determined using the Bioanalyzer 2100 (Agilent Technologies) using an RNA 6000 Nano kit. RNA quality was evaluated using the NanoDrop ND-1000 spectrophotometer (Thermo Scientific) and the Bioanalyzer 2100 (Agilent Technologies). A260/A280 and A230/A260 ratios greater than 1.8 and an RIN greater than 8 were considered acceptable. RNA samples were stored at −80°C. RNA sequencing (RNA-seq) was performed by BGI Tech Solutions (Copenhagen, Denmark) using non-stranded library preparation with mRNA enrichment (oligo(dT) magnetic beads), paired-end sequencing with 100 bp read length on the DNBSEQ platform.

### Bioinformatics

The paired-end raw sequence data had a total number of 7.99 E+08 reads (mean 6.66E+07 stdev 1.30E+07) and were quality controlled using FastQC (Galaxy Version 0.11.9). On average 78.2% (stdev 0.66%) of the reads could be uniquely mapped to the zebrafish reference genome GRCz11 using the RNAStar aligner (Galaxy Version 2.7.8a)^34^. The final transcript count data was generated using the HTSeq framework (Galaxy Version 0.9.1) for high throughput sequencing data^35^ based on Ensemble release 99 gene annotation using standard settings. All analysis was conducted on a private Galaxy instance running on the MedBioNode cluster at the Medical University of Graz. Further downstream analysis was conducted using R version 4.0.3 within the free RStudio Desktop version. Differential gene expression analysis was performed with DESeq2 package version 1.30.1^36^ on the count table as output from the HTSeq framework. DAVID Bioinformatics Resources 2021^37^ was used for pathway enrichment analysis by clustering DEGs and associated biological annotation terms into functional groups. The enrichment score cutoff in DAVID was set to 1.3, which corresponds to a corrected p value of 0.05.

### Behavioral screening

#### Impulsivity

The 5-CSRTT is a continuous performance test^38^ which has been extensively validated to measure impulsivity in zebrafish^33, 39^. Briefly, the fish (*n* = 9 – 10) is trained to respond regularly to a light stimulus in one of five spatially distinct locations on the rear wall of the test tank in order to gain a food reward (delivered at the front wall of the tank) in a purpose built testing arena (Zantiks AD, Cambridge, UK). Impulsivity is ascertained by examining the animal’s ability to withhold its response to the forthcoming light stimulus during a defined pre-stimulus interval (a variable-interval of 5-sec). In zebrafish, the test has been pharmacologically validated using atomoxetine, which reliably reduces impulsivity while not affecting other test parameters ^33, 39, 40^. The pretraining, and test phases, are described in detail in the **Supplementary Material**.

#### Risk-taking behavior

Novel object test Increased risk-taking behavior is common in ADHD patients^41^, and can be assessed in zebrafish using the novel object^42, 43^. We measured the time that zebrafish (*n* = 12 – 14 per group) spent close (within 2 cm) to a novel object (15 cm long black tube) in a novel tank (dimensions: 36 cm length × 27 cm height × 10 cm water column depth). Fish were recorded for 6 minutes and behavioral response was analysed using automated videotracking software (EthoVision, Noldus Information Technology Inc., Leesburg, VA - USA).

#### Hyperactivity

The open field test is commonly used for measuring locomotion and exploratory activity in adult zebrafish^44, 45^. Adult zebrafish (~4 months post-fertilisation: *n*= 35 habituation and *n*= 17 testing) were placed individually in a tank (20 cm length × 15 cm width, 10 cm water column depth) and filmed during 30-min exposure to an open field environment each day for 3 days. After 3 days of habituation, both *adgrl3.1^-/-^* and wild-type fish were placed individually in a beaker (300 mL) in home tank water or in a solution containing atomoxetine (*adgrl3.1^-/-^* only; 0.5 mg/L) for 30 min before being transferred to the open field test. All behaviors were analysed using automated video-tracking software (ANY-maze © – Stoelting Co., USA). The tank was separated into two virtual areas (central and peripheral area, 2 cm close to the wall) to provide a detailed evaluation of the exploratory activity. The following endpoints were measured: distance travelled (m), and immobility (s) Water was changed between each individual to minimise data variability^46^.

## Results

### *adgrl3.1^-/-^* increases externalising behaviors by altering noradrenergic signalling

Genetic variation in *Adgrl3* is associated with deficits in impulse control in rodents^26^ and *ADGRL3* polymorphisms (GWAS) have been linked to externalising, impulsivity-related disorders in humans ^12, 15^. We first investigated whether loss of *adgrl3.1* similarly reduced impulse control in adult zebrafish by using the 5-CSRTT (**Fig.1A**)^33, 39^. There was no difference in acquisition rates, nor any overall difference between WT and *adgrl3.1^-/-^* in the number of correct responses (**Fig. 1B**) during the pre-training stages of the 5-CSRTT. However, in the 5-CSRTT itself, compared to WT siblings, *adgrl3.1^-/-^* zebrafish displayed a significantly lower proportion of correct responses (**Fig. 1C**) reflecting inattention, and a greater number of anticipatory responses (**Fig. 1D**) reflecting impulsivity. There were a similar number of omissions in WT and *adgrl3.1^-/-^* (**Fig. 1E**) meaning that both genotypes completed the test the same number of times. The decreased attention and heightened impulsivity were more prominent in male *adgrl3.1^-/-^* zebrafish compared to females (**Supplementary Fig. 1A**). Treatment with atomoxetine partially reversed the attentional deficits and fully reversed the impulsivity in *adgrl3.1^-/-^* (**Fig. 1C,D**), suggesting that this gene modulates inattention (to a small degree), and impulsivity, predominantly via noradrenergic signalling.

**Figure 1.**
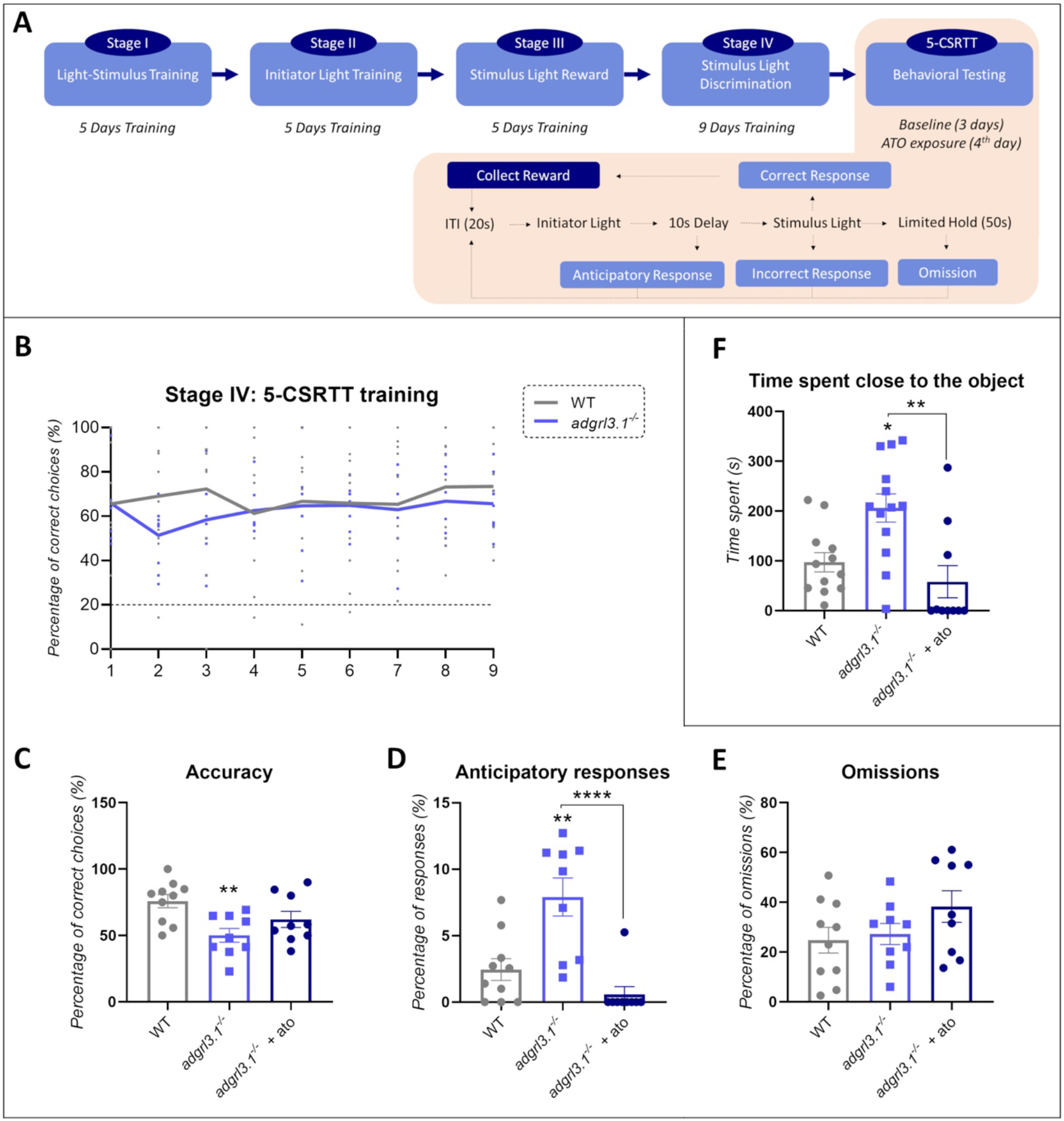
*adgrl3.1^-/-^* show attention deficits, impulsivity and increased risk-taking. **(A)** Flow-chart summarising the 5-CSRTT process. During the 5-CSRTT, fish were required to swim towards one of five spatially distinct LEDs when illuminated. Approaches to the illuminated light were ‘correct’ and the proportion of correct trials was a measure of attention. Prior to illumination, there was a variable-time (mean 5-sec) inter-trial interval, and responses during this interval were punished with subsequent non-reinforcement. Responses during this inter-trial interval (anticipatory or premature responses) were used as a measure of impulse-control. **(B)** No significant effects were found after two-way RM ANOVA for the acquisition during the Stage IV of the 5-CSRTT (Day*Group effect - F_(8, 136)_ = 0.50; *p* = 0.95; Group effect - F_(1, 17)_ = 0.69; *p* = 0.41; Day effect - F_(4, 247)_ = 0.45; *p* = 0.77). **(C)** One-way ANOVA yielded a significant effect for accuracy (F_(2,25)_ = 5.80; *p*\*\* = 0.0085), **(D)** anticipatory responses (F_(2,25)_ = 14.17; *p*\*\*\*\* < 0.0001) with no effects for **(E)** omissions (F_(2,25)_ = 1.80; *p* = 0.18). Tukey’s post-hoc analysis was used to characterise significant differences (*p*\*\* < 0.005 and *p* **** < 0.0001; *n* = 9 – 10). **(F)** Risk-taking behavior is defined as time spent close to the novel object. A significant ANOVA effect was observed for time spent close to the object (F_(2,32)_ = 8.35; *p*\*\* = 0.0012) where *adgrl3.1^-/-^* spent more time close to the object (*p*\*= 0.0148), an effect that was significantly decreased by atomoxetine in *adgrl3.1^-/-^* (*p*\*\* = 0.015; *n* = 12 – 13). The data is represented as mean ± S.E.M.

Children with ADHD and externalising personality dimensions have been also shown to have high levels of boldness in novel situations. This is strongly related to disruptive behaviors and conduct-related problems^47^. This behavior could be characterized as ‘high approach’ to unfamiliar situations, and represent a further facet of the ‘uncontrolled’ or impulsive phenotype^48^. For this reason, we next examined fish boldness in terms of their approach to a novel object^42, 43^. *adgrl3.1^-/-^* zebrafish spent more time close to the object (**Fig. 1F**), suggesting an increase in boldness. This increase was fully rescued by atomoxetine (**Fig. 1F**). There was no influence of sex effect on boldness in the novel object test (**Supplementary Fig. 1B**).

In summary, mutation of *adgrl3.1* made adult zebrafish significantly inattentive, impulsive and bolder, with a stronger effect in male animals for impulsivity (5-CSRTT) but not for boldness.

### *adgrl3.1^-/-^* zebrafish show noradrenaline-mediated hyperactivity after habituation to a novel environment

Previous studies with *adgrl3.1^-/-^* have focussed on single measures of hyperactivity as the endpoint^27, 28^. Here, we investigated the effect of loss of *adgrl3.1* function on motor activity over several test phases. Hyperactivity develops over time in children with ADHD (e.g.^49^). Similar patterns have been observed in *Adgrl3* knock-out rats^50^ as well as in spontaneously hyperactive rats [SHR]^51^, suggesting a strongly conserved mechanism. Therefore, prior to assessing hyperactivity, we first habituated the fish to their recording environment and analysed changes in their behavior. As predicted, we did not observe any differences in hyperactivity between WT and *adgrl3.1^-/-^* on the first three days of recording (**Fig. 2A**). However, *adgrl3.1^-/-^* showed a complex response pattern, spending significantly more time immobile than WT on day 1, an effect that reduced considerably on days 2 and 3 (**Fig. 2A**). This was consistent with the hypothesis that *adgrl3.1-/-* were experiencing higher anxiety on the first day^52^. On recording day 4 we found that *adgrl3.1-/-* were hyperactive compared to WT, swimming significantly further during the 5 minutes recording period. This phenotype was rescued by atomoxetine treatment, again suggesting a noradrenergic basis (**Fig. 2 B,C**). There was no difference in immobility between the genotypes at this timepoint (**Fig. 2C**). A sex effect was also observed for distance travelled: male *adgrl3.1^-/-^* showed the highest distance travelled and appeared to be driving the significant group differences (**Supplementary Fig. 1C**).

**Figure 2.**
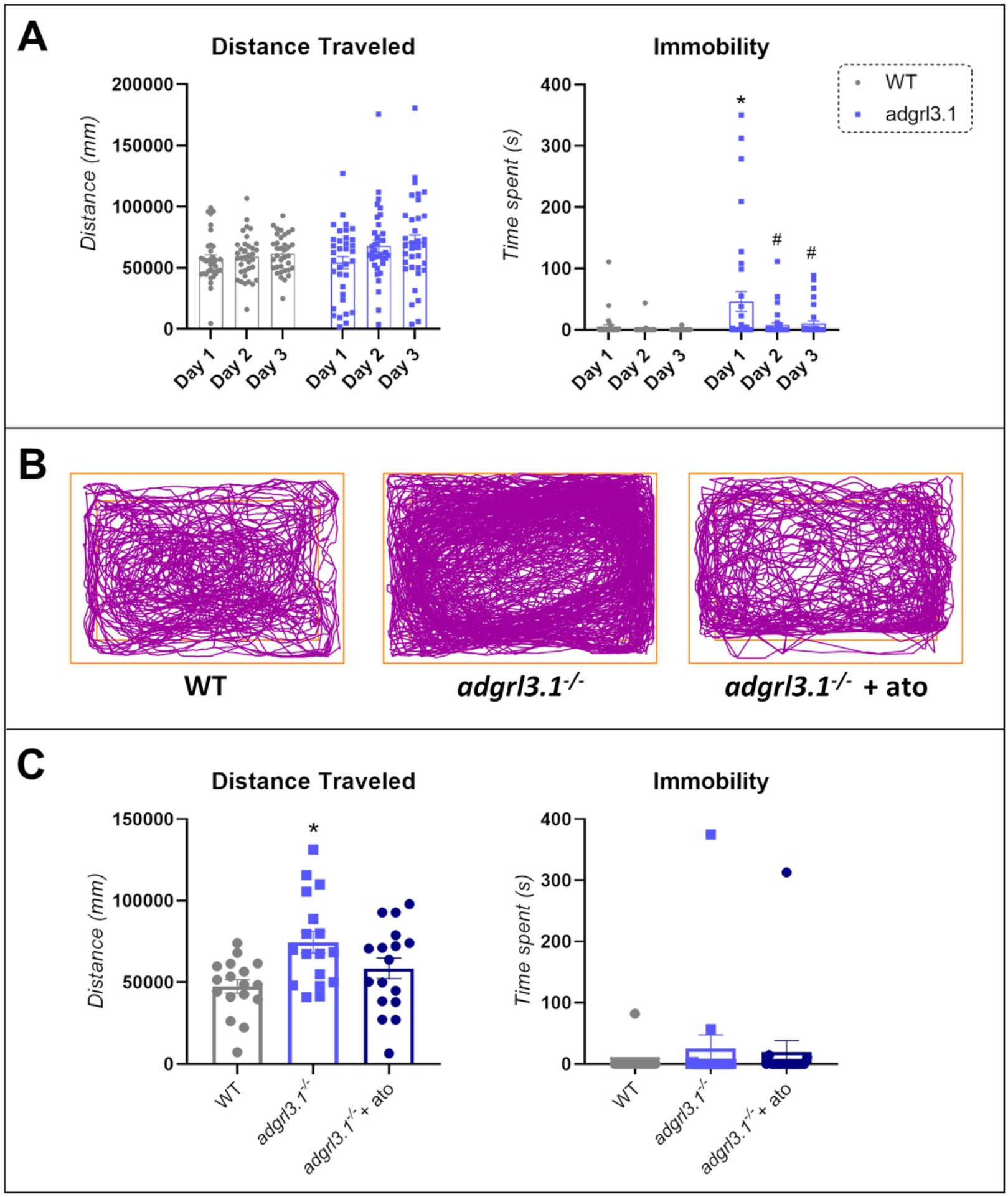
*adgrl3.1^-/-^* zebrafish show increased locomotion after 3 days of habituation. **(A)** A significant two-way RM ANOVA effect for habituation (F _(1.995, 135.7)_ = 3.84; *p*\* = 0.024) was observed for distance travelled, with no significant effects for interaction between factors (genotype*habituation; F _(2, 136)_ = 1.66; *p* = 0.19) nor genotype (F _(1, 68)_ = 1.40; *p* = 0.24). For immobility, a significant effect of genotype*habituation (F _(2, 136)_ = 3.66; *p*\* = 0.03), habituation (F _(1.998, 81.51)_ = 5.92; *p*\* = 0.013) and genotype (F _(1, 68)_ = 10.23; *p*\*\* = 0.002) was found. Post-hoc analyses showed that *adgrl3.1^-/-^* have increased immobility during the first day of habituation compared to WT (*p*\* = 0.018). *adgrl3.1^-/-^* also showed a habituation to the novel environment by showing a decrease in their immobility during the second day (*p*\* = 0.025) and third day (*p*\* = 0.03) compared to the first day of habituation. **(B)** Representative tracking of a WT *vs. adgrl3.1^-/-^ vs. adgrl3.1^-/-^* + ato animal during test day **(C)**A significant ANOVA effect was found for distance travelled (F_(2,48)_ = 5.44; *p*\*\* = 0.007) during the test day. Briefly, distance travelled was increased for *adgrl3.1-/-* compared to WT animals (*p*\*\* = 0.005) with no effect for *adgrl3.1^-/-^* compared to *adgrl3.1^-/-^* + ATO (*p =* 0.14). No effect for immobility was observed (*p* = 0.54). The data is represented as mean ± S.E.M.

### Genome-wide effects of *adgrl3.1* knockout identify novel functional pathways for treatment for EDs

#### Transcriptomic differences between the brains of *adgrl3.1^-/-^* and wild-type zebrafish

We next investigated transcriptomic differences between wild-type, *adgrl3.1^-/-^* and *adgrl3.1^-/-^* + ATO fish immersed acutely (20 min) in atomoxetine (*adgrl3.1^-/-^* + ATO; **Fig. 3**). Because atomoxetine rescued all the observed phenotypes, we examined acute treatment with atomoxetine to identify rapid effects of pharmacological alteration of noradrenergic signalling upon gene expression in *adgrl3.1^-/-^*. Principal component analysis (PCA) showed a clear separation between wild-type and *adgrl3.1^-/-^* (with and without ATO treatment) along PC1 of the plot explaining 54% of the total variance (**Fig. 3A**). In contrast, acute immersion in ATO did not lead to a large change in the transcriptome, with *adgrl3.1^-/-^* and *adgrl3.1^-/-^* + ATO clustering close together in the PCA plot. Differential expression analysis found a total of 869 differentially expressed genes between WT and *adgrl3.1^-/-^* and 896 differentially expressed genes between WT and *adgrl3.1^-/-^* + ATO (**Fig. 3B**). Only 34 genes were differentially expressed between *adgrl3.1^-/-^* and *adgrl3.1^-/-^* + ATO (**Fig. 3B**). Hierarchical clustering of the samples based on these differentially expressed genes showed consistent up- or downregulation of genes in all samples of a given group between WT and *adgrl3.1^-/-^* (**Fig. 3C**) and WT and *adgrl3.1^-/-^* + ATO (**Fig. 3E**). However, the few genes differing between *adgrl3.1^-/-^* and *adgrl3.1^-/-^* + ATO (**Fig. 3G**) showed considerable variation within the groups. We used volcano plots to highlight the most significant differentially expressed genes in each group. When comparing WT to *adgrl3.1^-/-^* we found heightened expression of genes including *atp6v1b2 (*ATPase H+ transporting V1 subunit *B2), birc6* (baculoviral IAP repeat containing 6) and *moesin a* (membrane-organizing extension spike protein a); and decreased expression of *dusp6* (dual-specificity phosphatase 6), *irs2a* (insulin receptor substrate 2a), *shisha4* (shisa family member 4), *bean1* (brain expressed, associated with NEDD4, 1) and *shisha7a* (shisa family member 7a) (**Fig. 3D**). A comparison of WT and *adgrl3.1^-/-^* + ATO revealed increased expression of the same genes as WT vs *adgrl3.1^-/-^*, as well as *nono* (non-POU domain containing, octamer-binding). The same genes showed decreased expression when comparing WT vs *adgrl3.1^-/-^* + ATO and WT vs *adgrl3.1^-/-^* **(Fig. 3F)**. Finally, when directly comparing *adgrl3.1^-/-^* to *adgrl3.1^-/-^* + ATO we saw a significant increase in *fosa* (v-fos FBJ murine osteosarcoma viral oncogene homolog a), *fosb* (v-fos FBJ murine osteosarcoma viral oncogene homolog b), *socs3a* (suppressor of cytokine signaling 3a), *carmil2* (capping protein regulator and myosin 1 linker 2) and *ripor2* (RHO family interacting cell polarization regulator 2), and a down-regulation of *birc2* (baculoviral IAP repeat containing 2), *rgs7b* (regulator of G protein signaling 7 binding protein b) and *pcf11* (PCF11 cleavage and polyadenylation factor subunit) **(Fig. 3H)**.

**Figure 3.**
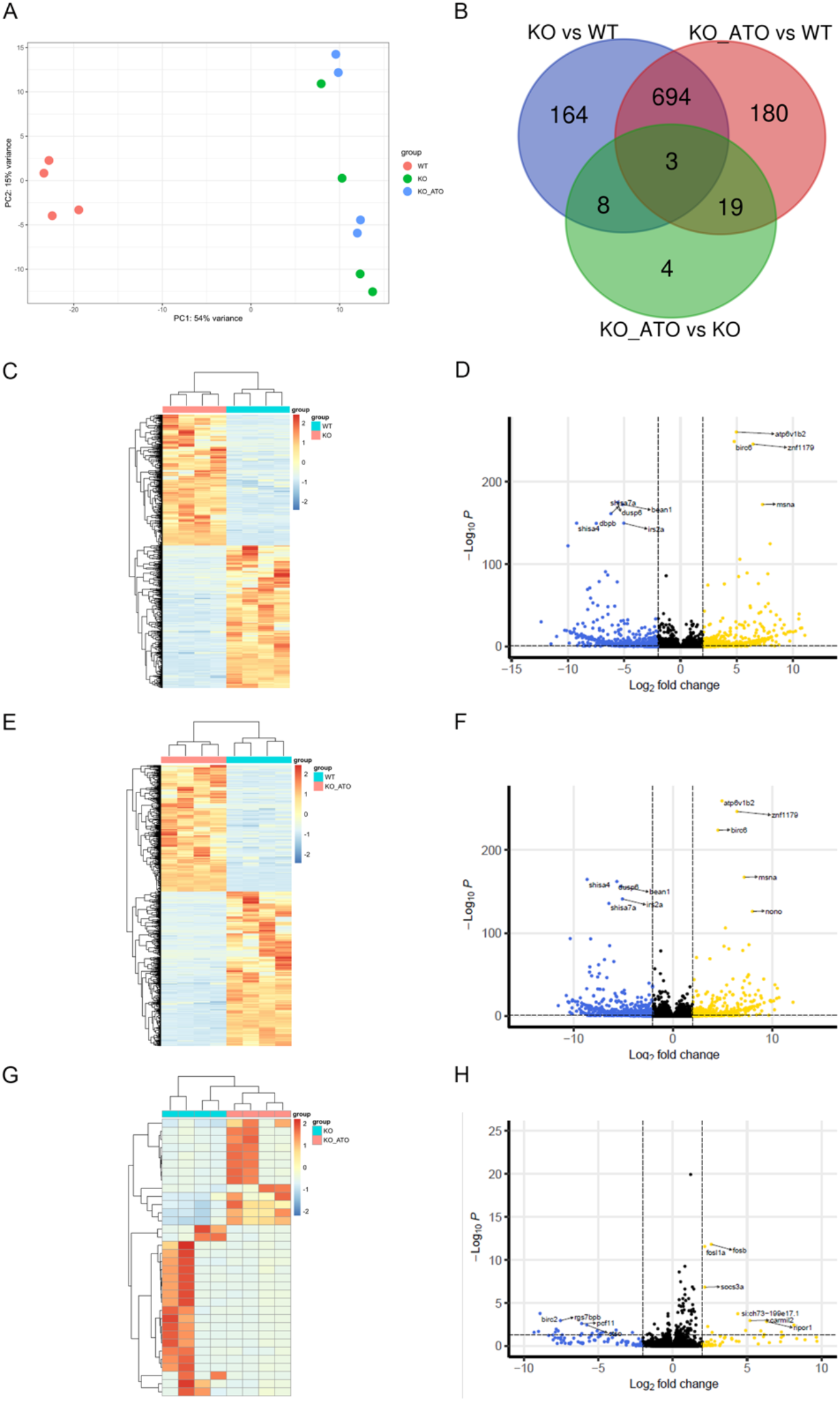
RNAseq summary data. **A**. Principal component analysis plot of the top 200 most variable genes after differential expression analysis. *n* = 4 brains/group. **B**. Venn diagram showing the overlap of differentially expressed genes (DEGs) between WT zebrafish, *adgrl3.1*^-/-^ and *adgrl3.1*^-/-^ treated with atomoxetine. **C**. Heatmap of DEGs (lowest adjusted *p* value (padj) < 0.05 and LFC > |2|) between WT and *adgrl3.1^-/-^*.**D**. Volcano plot displaying DEGs between WT and *adgrl3.1^-/-^*. DEGs with padj are highlighted. *n* = 4 per group. **E**. Heatmap of DEGs (padj < 0.05 and LFC > |2|) between WT and *adgrl3.1*^-/-^ treated with atomoxetine. **F**. Volcano plot displaying DEGs between WT and *adgrl3.1*-/- treated with atomoxetine. DEGs with padj are highlighted. *n* = 4 per group. **G**. Heatmap of DEGs (padj < 0.05 and LFC > |2|) between *adgrl3.1-/-* and *adgrl3.1*-/- treated with atomoxetine. **H**. Volcano plot displaying DEGs between *adgrl3.1*-/- and *adgrl3.1*-/- treated with atomoxetine. DEGs with padj are highlighted. *n* = 4 per group.

#### Pathway analysis reveals an enrichment of SPRY domain proteins and transcription factor activity

We used DAVID pathway^37^ analysis to investigate biological pathways underlying the observed gene expression changes and classify DEGS into functional clusters. We identified six significantly enriched clusters (enrichment > 2.43) in WT compared to *adgrl3.1^-/-^* (**Table 1**). The most enriched annotation cluster contained the GO term SPRY domain, followed by zinc-finger DNA binding domains and RNA polymerase II transcription factor activity in cluster 2. Other clusters included metabolism of drugs, xenobiotics and retinal (cluster 4) and immunoglobulins (cluster 5), suggesting that immune system activity may differ between WT and *adgrl3.1^-/-^*. Comparison of WT vs *adgrl3.1^-/-^* + ATO identified similar GO terms, including SPRY domain proteins in cluster 1, DNA binding and transcription factor activity in cluster 2, and lectins, interleukins and immunoglobulins in clusters 4, 5, 8 and 9 suggesting an important difference in immune system function between WT and *adgrl3.1^-/-^* (**Table 1**). In contrast, comparison between *adgrl3.1^-/-^* and *adgrl3.1^-/-^* + ATO did not reveal any enriched pathways.

**Table 1.** Enriched clusters and functional categories using DAVID pathway analysis in *adgrl3.1^-/-^* vs wild-type, and *adgrl3.1^-/-^* treated with atomoxetine vs wild-type.

| *adgrl3.1<sup>-/-</sup>* vs WT | *adgrl3.1<sup>-/-</sup>* + ATO vs WT |
| --- | --- |
| **Cluster 1 Score 26.89**<br>NACHT nucleoside triphosphatase<br>Domain: NACHT<br>SMO1288<br>Leucine-rich repeat<br>SPiA/Ryanodine receptor SPRY<br>Butyrophyrin-like<br>B30.2/SPRY domain<br>SPRY<br>DOMAIN: B30.2/SPRY domain<br>P-loop containing nucleoside triphosphate hydrolase<br>Intracellular signal transduction<br>PRY<br>Concanavalin A-like lectin/ glucanase subgroup | **Cluster 1 Score 25.05**<br>NACHT nucleoside triphosphatase<br>Domain: NACHT<br>SMO1288<br>Leucine-rich repeat<br>SPiA/Ryanodine receptor SPRY<br>Butyrophyrin-like<br>P-loop containing nucleoside triphosphate hydrolase<br>SPRY-associated<br>B30.2/SPRY domain<br>Intracellular signal transduction<br>SPRY<br>PRY<br>Concanavalin A-like lectin/ glucanase subgroup |
| **Cluster 2 score 5.7**<br>DOMAIN: C2H2-type<br>Zinc finger C2H2-type/integrase DNA-binding domain<br>ZnF-C2H2<br>RNA polymerase II transcription factor activity, sequence-specific DNA binding<br>Zinc finger<br>RNA polymerase II core promoter proximal region sequence-specific DNA binding<br>Zinc<br>Regulation of transcription from RNA polymerase II promoter<br>Metal-binding<br>Regulation of transcription, DNA-templated<br>Metal ion binding | **Cluster 2 Score 5.37**<br>DOMAIN: C2H2-type<br>Zinc finger C2H2-type/integrase DNA-binding domain<br>ZnF C2H2<br>RNA polymerase II transcription factor activity, sequence-specific DNA binding<br>RNA polymerase II regulatory region sequence-specific DNA binding<br>Zinc-finger<br>RNA polymerase II core promoter proximal region sequence-specific DNA binding<br>Regulation of transcription from RNA polymerase II promoter<br>Zinc<br>Metal-binding<br>Regulation of transcription, DNA-templated<br>Metal ion binding |
| **Cluster 3 Score 2.86**<br>Death-like domain<br>DAPIN domain<br>Domain: Pyrin<br>SMO1289 | **Cluster 3 Score 3.99**<br>Death-like domain<br>DAPIN domain<br>Domain: Pyrin<br>SMO1289 |
| **Cluster 4 Score 2.64**<br>Pentose and glucuronate interconversions<br>Drug metabolism – cytochrome P450<br>Retinal metabolism<br>Metabolism of xenobiotics by cytochrome P450<br>Steroid hormone biosynthesis<br>Drug metabolism – other enzymes<br>Porphyrin domain<br>Domain: UDPGT<br>glucuronosyltransferase activity<br>Ascorbate and aldarate metabolism<br>UDP-glucuronosyl/UDP-glucosyltransferase<br>UDP-glycosyltransferase activity<br>Glycosyltransferase<br>Biosynthesis of cofactors<br>Transferase activity, transferring glycosyl groups | **Cluster 4 Score 2.65**<br>hyaluronic acid binding<br>Link<br>DOMAIN: Link<br>LINK<br>C-type lectin fold<br>C-type lectin-like<br>Skeletal system development<br>Central nervous system development<br>Extracellular matrix<br>Cell adhesion<br>IGV |
| **Cluster 5 score 2.56**<br>Domain: IG<br>Immunoglobulin subtype<br>IG<br>Immunoglobulin-like fold<br>Immunoglobulin V-set<br>Domain: Ig-like<br>Immunoglobulin-like domain | **Cluster 5 Score 2.4**<br>CAAX prenyl protease 1<br>DOMAIN: Peptidase_M48_N<br>CAAX-box protein processing |
| **Cluster 6 Score 2.43**<br>CAAX prenyl protease 1<br>Domain: Peptidase M48 N<br>CAAX-box protein processing | **Cluster 6 Score 2.01**<br>Secreted<br>Extracellular region<br>Extracellular space |
|  | **Cluster 7 Score 1.76**<br>Immunoglobulin subtype<br>Immunoglobulin V-set<br>IG<br>Immunoglobulin-like fold<br>Immunoglobulin-like domain<br>DOMAIN: Ig-like |
|  | **Cluster 8 Score 1.38**<br>Necroptosis<br>C-type lectin receptor signalling pathway<br>Herpes simplex virus 1 infection<br>NOD-like receptor signalling pathway<br>Toll-like receptor signalling pathway |
|  | **Cluster 9 Score 1.34**<br>Interleukin-1 processing<br>Signalling by interleukins<br>Interleukin-1 family signalling |

## Discussion

In this study, we show that adult zebrafish lacking *adgrl3.1* displayed high levels of externalising behaviors (impaired impulse control and increased boldness in a novel situation), attentional deficits, and high levels of hyperactivity, all of which were rescued by atomoxetine. Our findings are thus consistent with the hypothesis that *ADGRL3* is associated with core aspects of externalising disorder phenotypes. We also found evidence of an increased stress response in a novel environment associated with *adgrl3.1^-/-^*, with high levels of immobility displayed during habituation to the recording setup prior to hyperactivity being expressed. Finally, differential gene expression (DEG) analysis comparing WT vs *adgrl3.1^-/-^* revealed several DEGs including *dusp6* (*MKP3*), which has been previously identified in a GWAS of ADHD patients^4^, and is known to modulate noradrenaline transporter (NET) activity. The DEG analysis also identified several putative functional pathways for externalising behaviors including enrichment of the SPRY domain, lending support to theories about the role of the immune system in ADHD and other EDs^53, 54^.

Disruption of *ADGRL3* function has been repeatedly linked to externalising symptoms in rodents^24, 55^ and zebrafish ^27, 28^. Here, we characterized several core externalising phenotypes in *adgrl3.1^-/-^*zebrafish, and the effect of atomoxetine on reducing these behaviors. *adgrl3.1^-/-^* performed a higher number of incorrect responses than WT in the 5-CSRTT suggesting attentional deficits. In the same task, *adgrl3.1^-/-^* showed a high level of impulsivity (higher anticipatory responses). Finally, *adgrl3.1^-/-^* displayed high rates of boldness in a novel environment, another core feature of externalising phenotypes^56^. Externalising behaviors such as impulsivity are debilitating and affect patients’ lives and outcomes. For example, an estimated 26.2% (95% CI: 22.7–29.6^57^) of prison inmates have ADHD. Increased externalising behaviors have been cited as key risk factors for criminality^58^ and predict the likelihood of delinquency^59^. Furthermore, other behaviors linked to higher rates of impulsivity (*e.g*. substance abuse) are more common in ADHD patients^12, 60^. *adgrl3.1^-/-^* has been linked to several ED, including substance abuse, and this has been shown to be independent of an ADHD diagnosis^12^. Furthermore, *ADGRL3* variants are associated with more extreme ADHD phenotypes (ADHD-combined; ADHD-C), worse outcomes in terms of disruptive behavior, persistence into adulthood and differential response to stimulant medication^61^. The fact that these behavioral phenotypes are so well conserved across vertebrate species, and even invertebrates^62^, strongly suggests that *ADGRL3* is functionally related to shared externalising phenotypes such as impulsivity and hyperactivity. This has significant implications for psychiatry, as dysfunctional *ADRGL3* may be implicated across a range of ED.

### Mechanisms of *adgrl3.1*-induced impulsivity and hyperactivity

Very little is known about the molecular function of *ADGRL3*, outside of its role in neuronal migration during development^63^. Here, for the first time, we found that the selective noradrenaline transporter inhibitor atomoxetine fully rescues several externalising phenotypes in *adgrl3.1^-/-^*, suggesting an important role for noradrenaline in ED. Previous work has linked *ADGRL3* variants to the dopamine system. Several markers of differential DA and NE activity have been identified in the striatum of *Adgrl3* knockout rats, including upregulation of presynaptic Tyrosine Hydroxylase and *Slc6a3* (which codes for the dopamine transporter), downregulation of *Dopamine receptor d1* in the striatum^24^, and upregulation of DAT in the prefrontal cortex^26^. These findings are conserved across vertebrates, with *adgrl3.1* morphant zebrafish displaying changes in the topography and number of dopaminergic neurons in the diencephalon^27, 28^. Collectively, these findings have led to the theory that *ADGRL3* exerts its behavioral effects via either direct or indirect effects on striatal dopamine^64^. Dysfunction of the DA system is an attractive hypothesis, given (for example) the therapeutic benefits of psychostimulant and non-psychostimulant medications in externalising disorders such as ADHD (e.g. atomoxetine blocks the presynaptic noradrenaline transporter, and thus can enhance both noradrenergic and dopaminergic signalling in the synapse). However, this is not necessarily useful from a clinical perspective, given the side effects of current ADHD medications, and the individual variability in therapeutic efficacy^2, 65^. Previous work has shown that transient knockdown of *adgrl3.1* in zebrafish larvae renders them more sensitive to stimulation of dopamine signalling^28^. This suggests that *adgrl3.1^-/-^* larvae have dysregulated NE levels, perhaps leading to saturation, as NE receptors regulate the response to dopamine agonists^66^.

To identify novel therapeutic targets for externalising symptoms, we carried out transcriptomic analysis of the whole brain of adult *adgrl3.1^-/-^* zebrafish and identified several DEGs. One notable DEG was *dusp6 (MKP3),* which has been previously identified in an ADHD GWAS^4^ and was downregulated in *adgrl3.1-/-*. Protein kinase c (PKC) regulates internalization of both DAT and NET, and recently *dusp6 (MKP3)*, a phosphatase that inactivates MAP kinases, has been shown to mediate PKC regulation of transporters^67, 68^. Therefore, it is possible that *dusp6* functions as a regulator of neuronal and synaptic plasticity^68^. Together with our data showing that atomoxetine rescues the behavioral phenotypes, this strongly implicates dysfunction of the NE system in more severe forms of ED.

*adgrl3.1* is a receptor for several ligands, including the Fibronectin leucine rich transmembrane protein 3 (*flrt3*)^69^. *adgrl3.1* is expressed presynaptically, and *flrt3* in the postsynaptic membrane, with interaction occurring via their extracellular tails. Deletion of the chromosome segment which includes *FLRT3* increases ADHD risk in humans, suggesting that loss of *FLRT3* function may cause the disease^70^. *flrt3* mediates cell-adhesion and sorting^71^ to control cell migration, axon guidance and axon outgrowth following injury, as well as activating Fibroblast growth factor (*fgf*) signalling^72^. FGF receptors are required for transcription of *dusp6*. Therefore, it is possible that the *fgf* pathway, which can activate target genes via MAPK/ERK^72^(a pathway which is known to modulate hyperactivity^73^) underpins externalising symptoms. *Adgrl3* and *Flrt3* have also been linked to synaptogenesis during development; antisense knock-down of *Adgrl3* or *Flrt3* in mouse reduces the number of glutamate synapses in the hippocampus^13^,^69^. However, the link between the dopaminergic phenotype of *adgrl3.1^-/-^* synaptogenesis is not clear. A subset of dopaminergic neurons co-release both dopamine and glutamate, and activation of dopamine D2 receptors inhibits both synapse formation and dopamine release^71^. Furthermore, ED-linked neural circuits may be fine-tuned by controlling the number of dopaminergic and glutamatergic synapses^70^. Therefore, it is possible that the noradrenergic phenotype of *adrgl3.1^-/-^* leads to a reduction of both dopamine and glutamate synapses, thus altering network activity and triggering ED symptoms. This may be an interesting target for future drug development.

The DAVID analysis revealed several pathways that were enriched in *adgrl3.1^-/-^*. Of particular interest was the SPRY domain, which was the most enriched cluster. The SPRY domain is involved with protein interactions in a diverse range of signalling pathways, ranging from innate immunity to RNA processing. Perhaps most interestingly, we found several enriched clusters relating to immune system function in *adgrl3.1^-/-^*. The links between externalising disorders and immune function are well established^74^. Children with ADHD have high levels of neuroinflammatory biomarkers such as Tumour Necrosis Factor-alpha (TNF-α)^74^. In addition, childhood atopic diseases (eczema, asthma) are associated with increased risk of ADHD^75^. Children with externalising behavioral problems have elevated levels of C-reactive protein (CRP) and interleukin 6 (IL-6)^76^. Despite this evidence for a link, cause and effect is hard to ascertain as studies are generally correlative in nature^77^. An example is studies into substance use disorders and neuroimmune function: these are confounded by the inflammatory response to administration of substances of abuse^78,79^. Here, we identified several pathways linked to innate neuroimmunity, suggesting potential shared genetic mechanisms.

Finally, although externalizing behaviors and EDs are consistently found to be more common in males than females, the biological basis of this difference is not clear^80, 81^. For example, gender differences are observed bidirectionally in impulsivity subtypes^80^ and it is hard to disentangle social and environmental factors that may impact on ED (e.g. normative gender roles and teacher/parental influences). Here, we found sex differences in attention, impulsivity and hyperactivity (both more prominent in male *adgrl3.1^-/-^* zebrafish compared to females), but not in novel object approach. This suggests some potential heritability of sex differences associated with these core ED phenotypes, and merits further investigation.

## Conclusion

In this study, we show that *adgrl3.1^-/-^* display strong, innate decreases in inhibitory control, increased risk-taking behavior, decreased attention and increased hyperactivity, suggesting an important genetic basis for externalising behaviors. All these behaviors were rescued by atomoxetine, suggesting a critical role of noradrenergic signalling in these phenotypes. Transcriptomic analyses of *adgrl3.1^-/-^* revealed several genes and pathways that may be useful for future study into the genetic basis of ED, and inform targets for future treatment.

## Conflict of Interest

The authors declare no conflict of interest.

## Acknowledgments

This study was financed in part by the Coordenação de Aperfeiçoamento de Pessoal de Nível Superior - Brazil (CAPES) - Finance Code 001 at the University of Portsmouth, UK. MOP currently receives funding from the NC3Rs (UK), and Dstl (UK). FR received funding from the Austrian Science Fund (grant numbers: J4090-B29 and P35774). The funders had no role in study design, data collection, and analysis, decision to publish, or preparation of the manuscript. We are grateful to Slave Trajanoski of the Centre for Medical Research at Medical University of Graz for biostatistical advice.

## Notes

### Competing Interest Statement

The authors have declared no competing interest.

