## Supplementary Material for "*adgrl3.1*-deficient zebrafish show noradrenaline-mediated externalizing behaviors, and altered expression of externalizing disorder-candidate genes, suggesting functional targets for treatment"

**Supplementary Methods**

**Animal handling and Randomization**

Housing conditions and water parameters are presented in the main file. Fish were fed three times/day with a mixture of live brine shrimp and flake food, except on the weekend where they were fed once/day. Fish used for RNA sequencing were removed from their housing tanks and immediately (controls) culled or exposed to atomoxetine (0.5 mg/L; Tokyo Chemical Industry (TCI)) for 30 min before culling, using the rapid cooling method. Brains were collected immediately. During the behavioral testing day, animals used in the new object boldness test and open field test (OFT) were either removed from their housing tanks and tested immediately, or treated for 30 minutes with atomoxetine, depending on the group. Animals used in the OFT were habituated to the test for 3 days (30 min habituation per day) before the exposure to atomoxetine and behavioral analysis of hyperactivity based on previous research using *Adgrl3* knockout rodents.

For the 5-CSRTT, animals were pair-housed for five weeks during the 5-choice training and then tested (as detailed in the 5-CSRTT protocol below). After behavioral testing, all fish were euthanized using 2-phenoxyethanol from Aqua-Sed (Aqua-Sed™, Vetark,Winchester, UK). All behavioral testing was carried out in a pseudo-randomized order, choosing fish at random from one of six tanks for each group. Fish were randomly pair housed and issued a subject ID, allowing all testing to be carried out in a fully blinded manner (i.e. experimenter and technical staff were blind to each animal’s genotype). Once all data was collected and screened for extreme outliers (*e.g.,* fish freezing, returning values of ‘0’ for behavioral parameters indicating non-engagement and high omission percentage (>95%)), the genotype was revealed, and data analyzed in full. No data were removed following unblinding. The experiments were carried out following scrutiny by the University of Portsmouth Animal Welfare and Ethical Review Board, and under license from the UK Home Office (Animals (Scientific Procedures) Act, 1986) [PPL: P9D87106F]. All behavioral tests were performed between 10 a.m. to 4 p.m. Finally, fish sex was defined by two experienced zebrafish researchers prior to behavioral testing, and confirmed by technical staff.

**5-CSRTT training phases and testing**

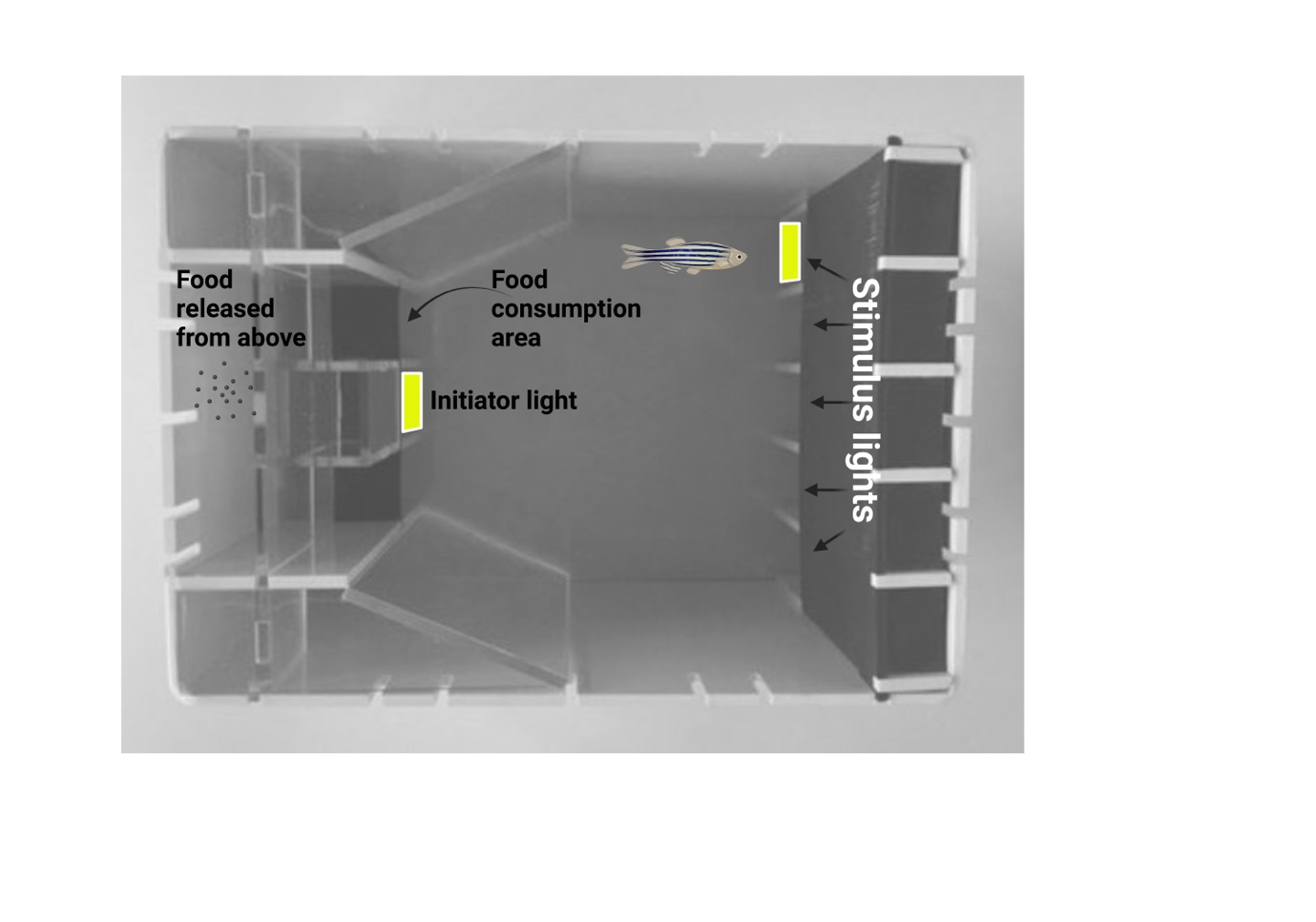

**Supplementary Fig 1 (S1).** 5-CSRTT training tank. During the testing period, to initiate a trial, the fish must swim into the ‘initiator light’ area. Once this is completed, the initiator light extinguishes, and a variable interval 10-sec delay is initiated. At the end of this interval, one of the five stimulus lights is illuminated (at random). If the fish makes a correct response (i.e. swims into the correct stimulus-light hole on the back wall), the stimulus light is extinguished, the food consumption area is illuminated, and food is released into the food hopper. Once the fish has consumed the food, the initiator light is illuminated ready for the next trial. Fish are trained on a sequence of stages to habituate them to the tank and to train stimulus-light responding.

Supplementary Fig 1 (S1) displays the 5-CSRTT equipment used in the study. The 5-CSRTT was performed, and data collected, using the Zantiks AD system (Zantiks Ltd., Cambridge, UK).

*1. Light training:* Fish were trained to trigger a response from the initiator and five stimulus lights in order to orient them to the stimuli for five days (one session of 30 trials/day). All lights were illuminated simultaneously in the initiator area and in the five openings at the back area of tank. The fish was required to enter in any of light area, so the light of the food consumption area, and the food reward is delivered via release from above the hopper. If animals do not respond in 60 seconds (limited hold), no food reward is delivered, and a new test begins after an interval of 20 seconds. The test is performed in five steps as described below.

*2. Initiator Light Training:* Here, fish learned to activate the initiator light (illuminated for 60 seconds). At this phase, when the initiator light is activated by the animal, it illuminates the feed hopper light, and food reward is delivered. In this phase, animals were trained for 5 days (one session of 30-trials/day) until the animal’s average of initiator light activation was 80% from 30 trials.

*3. Stimulus Light Reward:* In this phase, fish learned to approach any of the five illuminated stimulus lights to receive a food reward after triggering the initiator light (beginning test). Animals were trained for a minimum of 5 days, or until its average correct response rate was 70%, based on animals’ number of initiator light activation.

*4. Stimulus Light Discrimination:* Here, animals learn to discriminate between the individual stimulus lights. Once the fish activates the initiator light, one of the stimulus lights was activated (5-seconds delay). In subsequent trials, individual stimulus lights are illuminated in random order. The fish was required to approach the illuminated stimulus during a 50-sec illumination, and correct responses were rewarded with food delivery in the feed area. Animals were trained until they achieved a correct response rate of 60%, based on the animals’ total number of choices (min 9 days).

*5. 5-CSRTT:* The final stage was similar to Stage 4, where, following triggering the initiator light, one of five stimulus lights were illuminated (50-sec), and the fish was required to approach the correct light within the illumination time in order to receive a food reward. However, in this stage a variable-interval 10-seconds delay between activation of initiator light and the five-stimulus light is added to assess animals’ anticipatory responses. The parameters evaluated in 5-CSRTT were: number of correct responses (response to correct stimulus light), number of incorrect responses (incorrect location responses), omissions (does not approach stimulus), and premature responses (approach to any of the stimuli prior to illumination). These parameters were enabling the analysis of impulsiveness (premature responses), attention (correct responses) in adult zebrafish.

**Statistics**

Data were analyzed by using GraphPad Prism and the results were expressed as means ± standard error of the mean (S.E.M). Normality was assessed *a priori* and samples with normal distribution were tested as follows. To assess whether there were any effects of genotype or for *adgrl3.1^-/-^* treated with atomoxetine during the OPT, 5-CSRTT and risk-taking behavior, a one-way ANOVA (three groups - WT *vs. adgrl3.1^-/-^ vs. adgrl3.1^-/-^* + ato) was used followed by Tukey’s post-hoc analysis. Two-way ANOVA followed by Tukey’s post-hoc analysis was used to analyze sex effects (genotype*sex) in the OPT, risk-taking behavior and 5-CSRTT, and for habituation analysis (genotype*days). Results were considered significant when p ≤ 0.05.

**Supplementary Results**

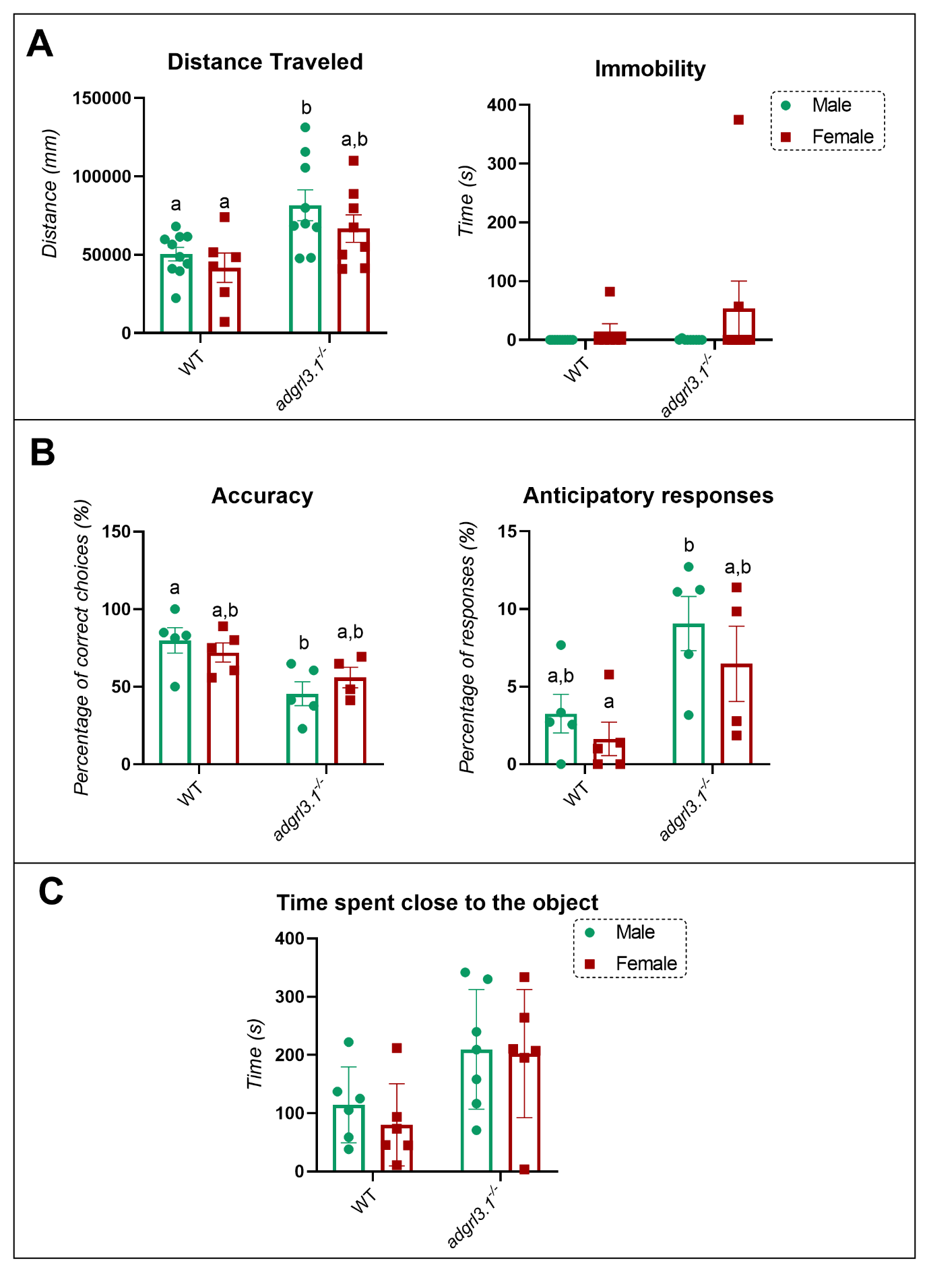

**Supplementary Fig 2.** Sex effect for *adgrl3.1^-/-^* mutant compared to WT animals in the open field, 5-CSRTT, and novel object test.

**Supplementary figure 2 (S2)** displays the sex differences we observed during behavioral protocols. During the locomotor pattern analysis (Figure S2A), a significant effect for sex was found in the distance travelled (F _(1, 29)_ = 11.69; *p*** = 0.0019) with no interaction between factors (F _(1, 29)_ = 0.1415; *p* = 0.7096) neither an effect for genotype (F _(1, 29)_ = 2.057; *p* = 0.1622). Interestingly, an increase in distance travelled was observed only for male *adgrl3.1^-/-^* when comparing to males WT (*p* * = 0.0315) with no effect comparing females *adgrl3.1^-/-^* and females WT (*p* = 0.2121). No significant differences were observed for immobility when looking at the effect for interaction between factors (F _(1, 29)_ = 0.7291; *p* = 0.4002), genotype (F _(1, 29)_ = 0.7562; *p* = 0.3917), and sex (F _(1, 29)_ = 2.070; *p* = 0.1609). Tukey’s test was used as post-hoc analysis and different letters indicate significant statistical differences (*p* < 0.05; *n =* 6 – 10). Figure **S2B** displays the effects of sex in the 5-CSRTT. Although no interaction (genotype *vs.* sex) or sex effect was observed for the parameters studied here, a significant genotype effect was observed for accuracy (F _(1, 15)_ = 11.84; *p*** = 0.0036) and anticipatory responses (F _(1, 21)_ = 10.78; *p*** = 0.0050). Tukey’s test was used as post-hoc analysis and different letters indicate significant statistical differences (*p* < 0.05; *n =* 4 – 5). **Finally C)** Similarly, only a genotype effect was observed for time spent close to the new object (F _(1, 21)_ = 9.076; *p*** = 0.0066; *n =* 6 – 7). The data is represented as mean ± S.E.M.
